# Universal antibiotic tolerance arising from antibiotic-triggered accumulation of redox metabolites

**DOI:** 10.1101/453290

**Authors:** Kui Zhu, Shang Chen, Tatyana A. Sysoeva, Lingchong You

## Abstract

*Pseudomonas aeruginosa* is an opportunistic pathogen that often infects open wounds or patients with cystic fibrosis. Once established, *P. aeruginosa* infections are notoriously difficult to eradicate. This difficulty is in part due to the ability of *P. aeruginosa* to tolerate antibiotic treatment at the individual-cell level or through collective behaviors. Here we describe a new mechanism by which *P. aeruginosa* tolerates antibiotic treatment by modulating its global cellular metabolism. In particular, treatment of *P. aeruginosa* with sublethal concentrations of antibiotics covering all major classes promoted accumulation of the redox-sensitive phenazine - pyocyanin (PYO). PYO in turn conferred general tolerance against diverse antibiotics for both *P. aeruginosa* and other Gram-negative and Gram-positive bacteria. We show that PYO promotes energy generation to enhance the activity of efflux pumps, leading to enhanced antibiotic tolerance. This property is shared by other redox-active phenazines produced by *P. aeruginosa*. Our discovery sheds new insights into the physiological functions of phenazines and has implications for designing effective antibiotic treatment protocols.

**Author Summary:** Antibiotic tolerance can facilitate the evolution of resistance, and here we describe a previously unknown mechanism of collective antibiotic tolerance in *Pseudomonas aeruginosa*. In particular, *P. aeruginosa* treated with sublethal concentrations of antibiotics covering all major classes promotes accumulation of pyocyanin (PYO), an important virulence factor. In turn, PYO confers general tolerance against diverse antibiotics for both *P. aeruginosa* and other bacteria. Our discovery is a perfect example of what Nietzsche once said: *That which does not kill me makes me stronger*.

## Introduction

The overuse and misuse of antibiotics have led to a global crisis (1): bacteria have developed resistance against every existing antibiotic and are doing so at an alarming rate, considering the timescale at which new antibiotics progress from development to clinical application (2, 3). The drying antibiotic pipeline further heightens the global threat created by infectious bacteria (4, 5). A critical approach to the problem is developing ways to revitalize existing antibiotics (6). To extend the use of existing antibiotics, we need to develop a mechanistic understanding of the diverse ways by which bacteria survive antibiotics. Such an understanding is critical for designing therapeutic approaches that subvert the survival tactics by bacterial pathogens.

Past efforts on antibiotic resistance have focused on responses of individual cells, such as mutations in the antibiotic targets, enzymatic activity that inactivates antibiotics, and increased activation of efflux pumps (6). Unlike antibiotic resistance, which is due to inherited or acquired mutations (7, 8), tolerance reflects the ability of individual cells (9) or cell populations (10) to survive antibiotic treatment without acquiring new mutations. However, it has been long realized that antibiotic tolerance precedes resistance (11–13). Recent *in vitro* experiments have shown that antibiotic tolerance can facilitate the evolution of resistance (14). In particular, antibiotic tolerance can pave the way for the rapid subsequent emergence of antibiotic resistance in bacteria; thus, preventing the evolution of tolerance may shed light on alternative strategies for antibiotic treatments. Importantly, the ability to survive antibiotic treatment is typically considered an intrinsic property of the single bacterial cells or bacterial populations, before antibiotics are applied. In contrast to these prevailing views, here we describe a previously unknown mechanism of collective antibiotic tolerance in *Pseudomonas aeruginosa*.

PYO, one of the most studied phenazines, is a redox-active metabolite giving the characteristic blue-green pigment of *P. aeruginosa* cultures (15). PYO is typically produced when *P. aeruginosa* enters the stationary phase when the cell density is high (16). Recent studies have demonstrated a number of physiological roles of PYO, such as serving as a signaling compound (17), facilitating biofilm development (18), promoting iron acquisition (19), and influencing colony formation (20). It is also known to confer a broad-spectrum antibiotic activity (21). Our work provides novel insights into the biological function of bacterial redox-active metabolites, and suggests a role of such metabolites, including PYO and other phenazines, in the survival of bacteria during acute stress.

## Results

### Subinhibitory concentrations of antibiotics induce PYO accumulation

We observed that sub-inhibitory concentrations of kanamycin (Kan), a commonly used aminoglycoside antibiotic, induced a blue-greenish color change in *P. aeruginosa* PAO1 (ATCC 47085) cultures. PYO is a typical pigment for such color change (15). We confirmed this notion by purifying PYO using an established method (22), which also allowed us to measure PYO accumulation. The induction of PYO followed a biphasic dependence on the Kan dose. The blue-green color of the cultures became more visible with increasing Kan concentrations (**Fig. 1 *A***). This color change is consistent with direct quantification of PYO in the supernatants (**Fig. 1*B***). Concurrently, the bacterial density (A600 nm) decreased with the Kan concentrations (**Fig. 1*B***). This enhanced PYO accumulation related to the culture without antibiotic treatment was transient, as evident in the direct measurements of PYO purified from the supernatants of cultures treated with 20 μg/mL Kan (**Fig. 1 *C***).

**Fig. 1.**
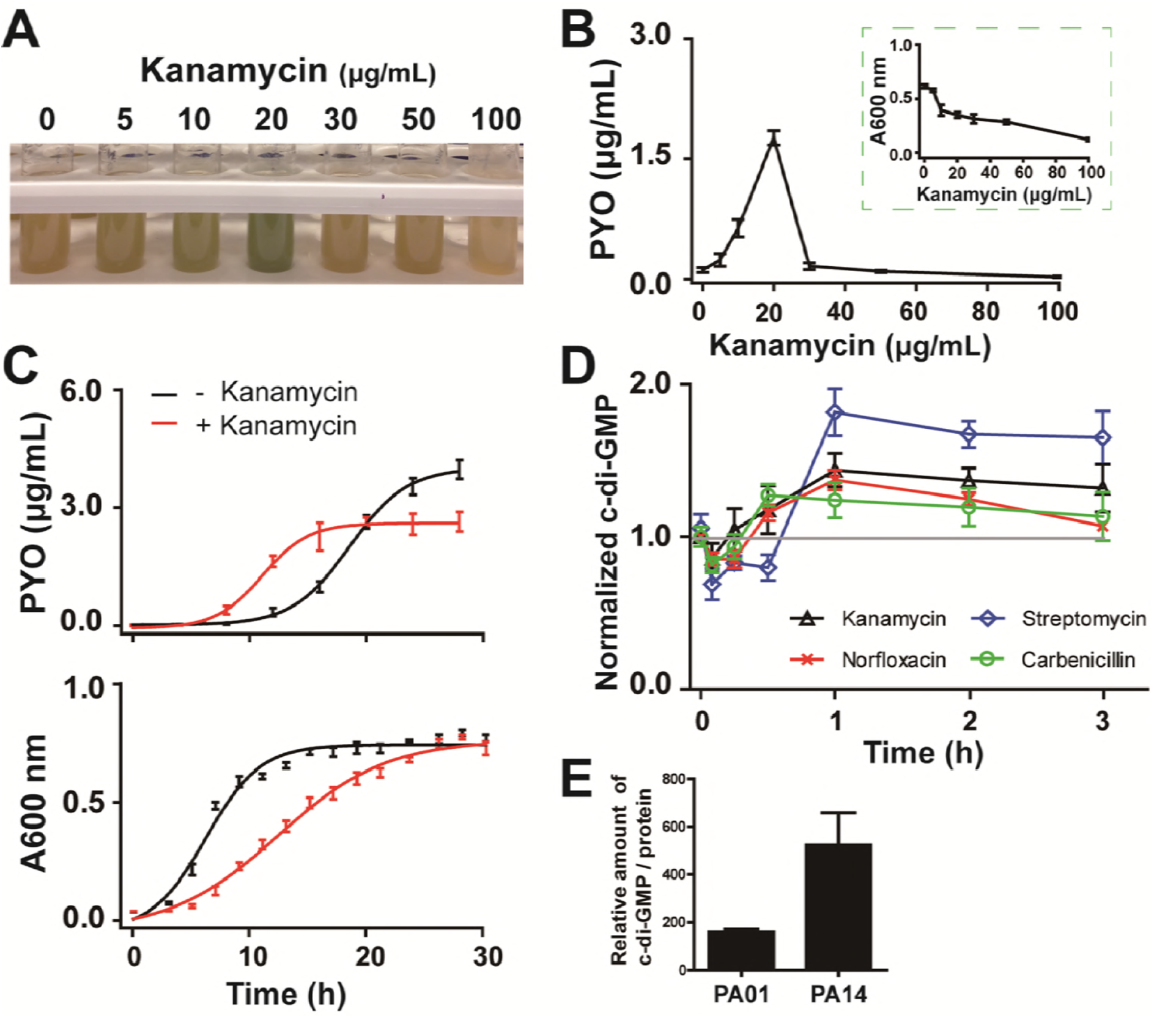
**Antibiotics induced accumulation of PYO**. (**A**) Kanamycin induced color changes in *P. aeruginosa* PA*O1* cultured in LB medium. (**B**) PYO concentration and cell density (A600 nm) of PAO1 in varying concentrations of kanamycin. (**C**) Typical dynamics of PYO concentrations and bacterial growth curves of PAO1 treated with 20 μg/mL kanamycin. (**D**) Different classes of antibiotics promoted c-di-GMP accumulation in PAO1. Each data point was normalized with respect to the concentration of c-di-GMP in the absence of antibiotic treatment. Means ± s.d. were presented for **A** to **C** (n = 3), for **D** (n = 6).

We wondered whether the enhanced PYO accumulation could represent a general stress response to antibiotic treatment. To test this notion, we measured PAO1 strain responses to other antibiotics with different modes of action, including chloramphenicol that targets ribosome 50S subunit, norfloxacin (quinolone family) that inhibits DNA replication, polymyxin B (polypeptide family) that alters cell membrane permeability, and carbenicillin (β-lactams) that inhibits cell-wall synthesis. All tested antibiotics promoted accumulation of PYO in a similar biphasic manner (**Supplementary, Fig. S1 *A***). This response was not unique to PAO1. The PA14 strain, which is known to produce more PYO than PAO1 (17), exhibited qualitatively the same responses to these antibiotics (**Supplementary, Fig. S1 *B***). The biphasic PYO accumulation and its temporal dynamics reconcile the apparently contradictory conclusions on PYO accumulation under antibiotic stresses measured previously by single-point measurements (23–25).

Cyclic di-GMP (c-di-GMP), an intracellular second messenger, has been shown to confer tolerance to antibiotics (26, 27). Moreover, c-di-GMP has been shown to promote PYO production (28). These studies suggest a potential role of c-di-GMP in antibiotic-mediated PYO accumulation. Consistent with this notion, we observed increased levels of c-di-GMP in PAO1 in response to several antibiotics (**Fig. 1*D***). In the absence of antibiotics, the accumulation of c-di-GMP was approximately 2.5 fold higher in PA14 than in PAO1 during early stationary phase (**Supplementary, Fig. *S2A***). This observation provides a potential explanation for the typically higher PYO accumulation in PA14 than in PAO1. Additionally, exogenous GTP – the precursor of c-di-GMP (27), leads to increased c-di-GMP over time and increased PYO accumulation in a dose-dependent manner (**Supplementary, Fig. S2*B*, S2*C***, and **S2*D***). This result confirms the recent finding that PYO production is c-di-GMP dependent (29).

### Accumulated PYO enhances antibiotic tolerance in bacteria

We wondered whether the enhanced accumulation of PYO could represent a survival mechanism for the population under acute antibiotic stress. Indeed, growth of *P. aeruginosa* PAO1 in the presence of Kan was enhanced by exogenously added PYO (**Fig. 2*A***). This PYO-mediated tolerance was effective against other aminoglycosides (gentamicin, streptomycin and tobramycin) and antibiotics of other classes (norfloxacin, chloramphenicol and carbenicillin) (**Fig. 2*A***). In the presence of PYO, PAO1 cultures exhibited a much shorter lag time before recovering in the presence of an antibiotic, in comparison to cultures without exogenously added PYO (**Supplementary, Fig. S3**). Endogenous PYO, if accumulated at a sufficiently high level such as in the cystic fibrosis respiratory tract (30), can also provide protection. In addition, PYO-mediated tolerance was maintained when PAO1 was cultured in other media or different oxygen conditions (**Supplementary, Fig. S4**). An exception to the general PYO-mediated tolerance was polymyxin B, where PYO enhanced its ability to inhibit bacterial growth (**Fig. 2*A***).

**Fig. 2.**
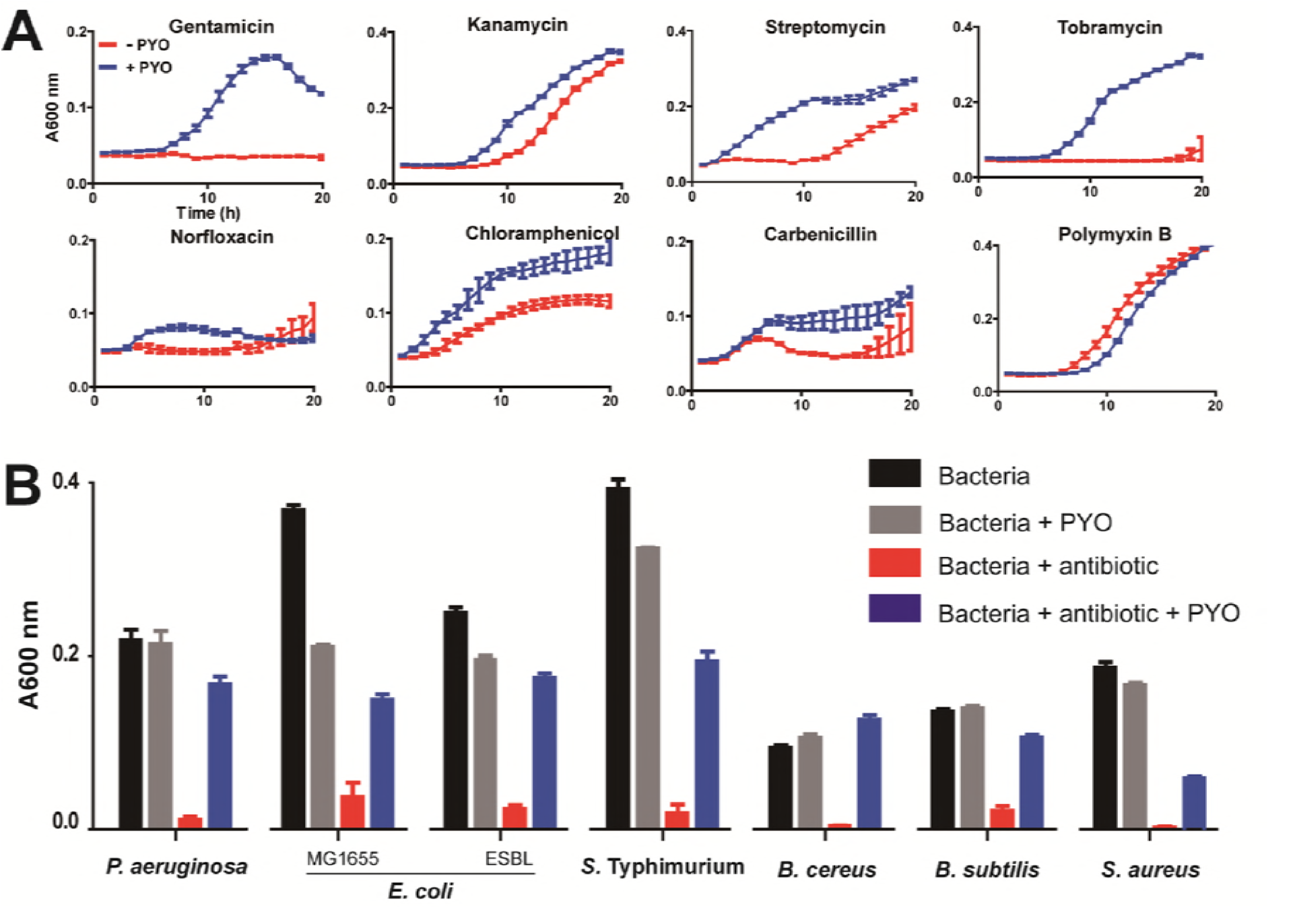
**PYO-mediated tolerance against antibiotic treatment**. (**A**) PYO-mediated tolerance in *P. aeruginosa* PAO1. Various antibiotics were tested, including 50 μg/mL kanamycin, 4 μg/mL gentamicin, 20 μg/mL streptomycin, 20 μg/mL tobramycin, 0.5 μg/mL norfloxacin, 7.5 μg/mL chloramphenicol and 50 μg/mL carbenicillin and 1.0 μg/mL polymyxin B. Means ± s.d. were presented (n = 3). (**B**) PYO-mediated tolerance in Gram-negative bacteria, including *P. aeruginosa* PAO1, *Escherichia coli* (MG1655 and ESBL-expressing strain), and *Salmonella* Typhimurium, and Gram-positive bacteria, including *Bacillus cereus, Bacillus subtilis*, and *Staphylococcus aureus*. Bacteria were cultured in LB media supplemented with 20 μg/mL streptomycin, except 5 μg/mL streptomycin for *B. cereus*. Means ± s.d. were presented (n = 6). *p*-values between streptomycin treated bacteria with and without PYO were less than 0.01. In addition to PAO1, PYO conferred antibiotic tolerance to several other strains of *P. aeruginosa*, including PA1C (ATCC 15692), PAO-JP2 with *ΔlasI* and *ΔrhlI* (31), PAO1-W and its mutants (PAO-mxM and PAO-mxS) (32), PA14 and its mutant (PA14 *Δphz*) (17). Among these, PAO-mxM, PAO-mxS and PA14 *Δphz* cannot synthesize PYO, indicating that PYO-mediated tolerance is independent of the strain’s ability to synthesize it (**Supplementary, Fig. S5**). The tolerance was maintained for other Gram-negative *(Escherichia coli, Salmonella* Typhimurium) and Gram-positive *(Bacillus cereus, Bacillus subtilis* and *Staphylococcus aureus)* bacteria treated with antibiotics (**Fig. *2B***).

### Oxidized PYO serves as electron acceptors to mediate antibiotic tolerance

As the PYO-mediated tolerance was general against different classes of antibiotics for diverse bacteria, it likely resulted from PYO-mediated modulation of cellular physiology that is common for all these bacteria. As a redox-active molecule (33), PYO can exist in reduced or oxidized state, depending on its chemical environment. The oxidized form of PYO can act as an electron acceptor to modulate the global energy metabolism in the cell by regulating the flux of electron shuttling (34). Similar functions have been proposed for other chemicals that can act as electron acceptors. For instance, the electron-shuttling ability of oxidized tetrathionate (35) and nitrate (36) has been implicated in enhancing bacterial growth advantage by promoting respiration. Additionally, hydrogen sulfide has been shown to enhance bacterial survival during antibiotic stress (37). Consistent with this notion, several reductants, including nicotinamide adenine dinucleotide (NADH), β-mercaptoethanol, and *N*-acetyl-*L*-cysteine (NAC), all eliminated PYO-mediated antibiotic tolerance (**Fig. 3*A*** and **Supplementary, S6*A***). By themselves, these chemicals did not affect *P. aeruginosa* growth or response to antibiotics. The presence of these reductants in excess prevented significant accumulation of the oxidized PYO, thus interfering with its ability to act as an electron acceptor. As a side, our results might provide an explanation to previously observed unknown benefit of reductants as antibiotic-adjuvants in treating CF patients (38), and suggest new strategy to devise antibiotic adjuvants.

**Fig. 3.**
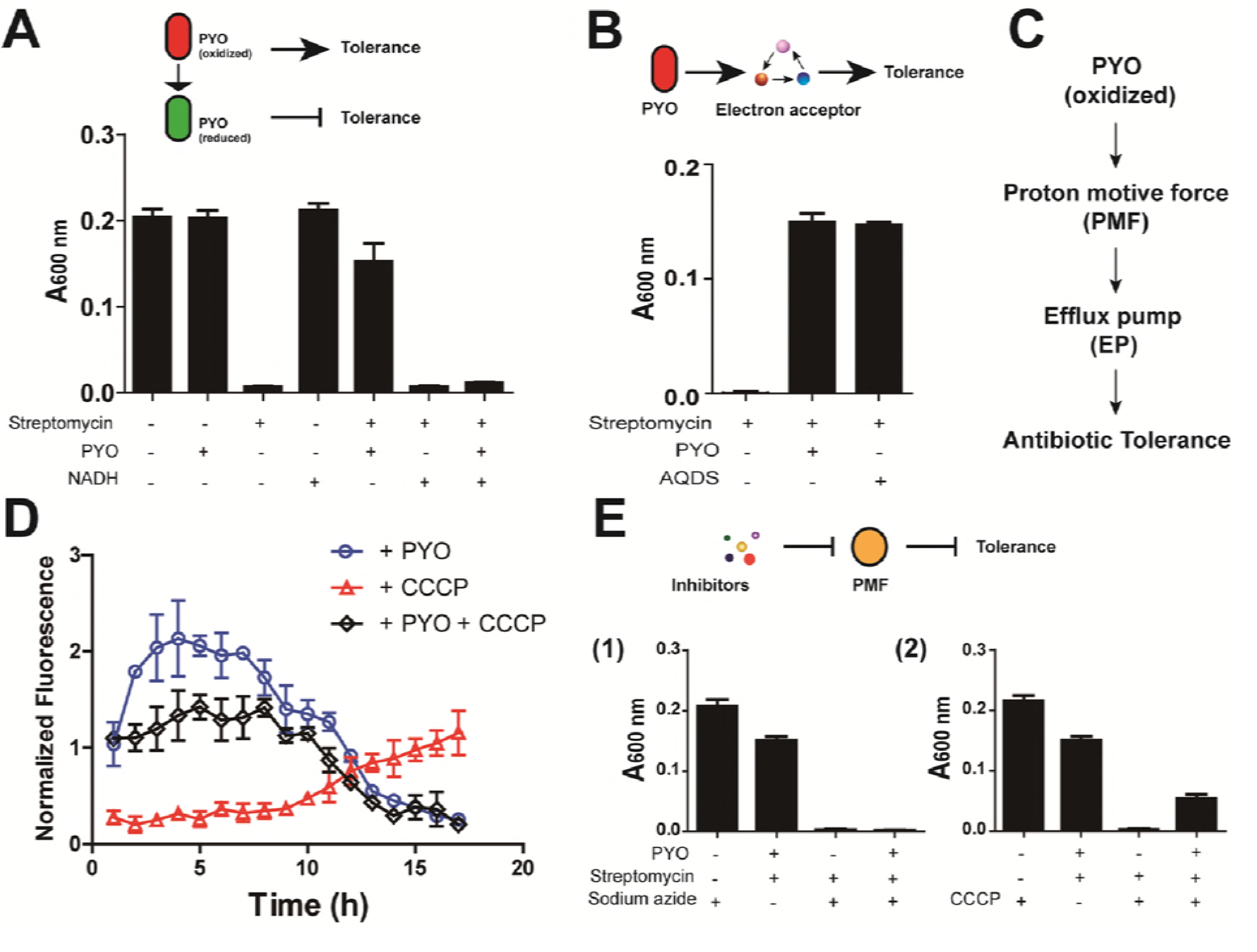
**PYO mediates antibiotic tolerance by enhancing the cellular metabolism**. (**A**) PYO-mediated tolerance depends on the oxidized form of PYO. Cultures of *P. aeruginosa PAO1* treated with 20 μg/mL streptomycin was supplemented with 15 μmol/L NADH and 2 μg/mL PYO. (**B**) PYO served as an electron acceptor. AQDS (1 μg/mL) is a compound that serves as an electron receptor. It shows similar effect as PYO in offering protection of *P. aeruginosa* PAO1 against streptomycin treatment. (**C**) Scheme of PYO-mediated tolerance. ATP, adenosine triphosphate; EP, efflux pump; NAD+, oxidized form of nicotinamide adenine dinucleotide; NADH, reduced form of nicotinamide adenine dinucleotide; PMF, proton-motive force; WST-8, water-soluble tetrazolium salt. (**D**) PYO promoted proton-motive force (PMF). Representative membrane potential of 20 μg/mL streptomycin treated *P. aeruginosa* PAO1 in the presence of either 2 μg/mL PYO or 10 μmol/L CCCP, or both as indicated. Membrane stain (DiOC_2_(3)) was simultaneously added to the cultures to final concentration of 100 μmol/L with streptomycin and PAO1, for long term fluorescent measurement. Fluorescence of streptomycin treated PAO1 in the presence of either PYO or CCCP, or both, was normalized to that of streptomycin treated PAO1. (**E**) PYO-mediated tolerance was PMF dependent. Dissipation of PMF by inhibitors such as sodium azide and CCCP could significantly decrease the tolerance. Means ± s.d. were presented for **A** to **C**, and **E** (n = 6), for **D** (n = 9).

If electron-shuttling ability is critical for PYO-mediated antibiotic tolerance, we reasoned that other redox-active phenazines might exhibit similar protective effects. Indeed, several natural analogs of PYO, including phenazine-1-carboxylic acid (PCA), phenazine-1-carboxamide (PCN) and 1-hydroxyphenazine (1-OHPHZ), promoted *P. aeruginosa* survival during antibiotic exposure (**Supplementary, Fig. S6*B***). Further confirming this notion, two other well-established electron acceptors, 2,6-anthraquinone disulfonate (AQDS) (39), and methylene blue (MB) (40), also conferred antibiotic tolerance in *P. aeruginosa* (**Fig. 3*B*, 3*C*** and **Supplementary, S6*B***).

### Oxidized PYO stimulates energy metabolism to pump antibiotics out

Studies have suggested a role of PYO in promoting ATP production (34), particularly when cells enter stationary phase or are under stress (40). Consistent with this notion, we observed that exogenously added PYO promoted ATP accumulation during stationary phase (**Supplementary, Fig. S7**). An increase in ATP accumulation can facilitate generation of the proton-motive force (PMF) (41), which plays a critical role in promoting bacterial survival (but not growth) under anaerobic condition (34), in cell division (42) and increased efficiency of efflux pumps (EPs) (43, 44). To test this notion, we measured the membrane potential with the carbocyanine dye DiOC_2_(3), which confirmed an elevated PMF in the presence of PYO (**Fig. *3D***). Additionally, the multidrug resistance phenotype in *P. aeruginosa* is in major part due to the expression of abundant EP systems (45, 46). For example, the MexAB-OprM pump plays a key role in the intrinsic resistance for several types of antibiotics (44). An increasing efficiency of EPs, mediated by an elevated PMF, would provide an explanation for the general PYO-mediated tolerance to a wide variety of antibiotics. The PYO-mediated promotion of PMF also explains the lack of PYO-mediated tolerance against polymyxin B (**Fig. *2A*** and **Supplementary, *S3B***): the loss of cell membrane integrity caused by polymyxin B completely eliminates the PMF. This is consistent with the previous observation that none of EPs extrude polymyxin B in *P. aeruginosa* (47), because EPs are anchored in membrane and most are driven by PMF (43–45). Aside from polymyxin B, an increased PMF may not always benefit the cells. In particular, as the uptake of aminoglycosides is energy dependent, an increased PMF can enhance their uptake and subsequent inhibition of bacterial growth (48) (i.e. by overcoming the increased efficiency of EPs). Thus, we reasoned that PYO-mediated tolerance would be eliminated at high levels of aminoglycosides. Indeed, 100 μg/mL Kan and 10 μg/mL gentamycin effectively suppressed *P. aeruginosa* growth, further demonstrating that PYO mediated tolerance was due to PMF.

Given this model (**Fig. 3*C***), we expected that inhibition of the hydrolysis of ATP would abolish PYO-mediated antibiotic tolerance. Indeed, sodium azide, which inhibits the F_1_ subunit of ATP synthase (49), abolished the tolerance (**Fig. *3E***). Similarly, *N,N’*-dicyclohexylcarbodiimide (DCCD), which blocks the proton translocation channel in the F_0_ subunit of ATP synthase (34), also reduced the PYO-mediated tolerance (**Supplementary, Fig. S8 *A***). Further confirming the role of PMF, ionopore carbonyl cyanide *m*-chlorophenyl hydrazone (CCCP), which directly dissipates PMF, also reduced PYO-mediated tolerance (**Fig. *3E***). Furthermore, both CCCP and sodium azide enhanced growth inhibition by antibiotics, even in the presence of PYO (**Supplementary, Fig. S8*B*** and **S8*C***).

We further tested modulation of EP efficiency by quantifying intracellular accumulation of ethidium bromide (EtBr), a well-established substrate of EPs in *P. aeruginosa* (44). Indeed, PYO accelerated the extrusion of EtBr by cells (**Fig. *4A***). If an enhanced efficiency of EPs was responsible for PYO-mediated antibiotic tolerance, we would expect chemical inhibitors of EPs to abolish this tolerance. We tested this notion by using phenylalanine-arginine β-naphthylamide (PAβN), which inhibits the resistance nodulation cell division (RND) pumps, the major contributors to antibiotic resistance in *P. aeruginosa* (50). Indeed, PAβN reduced PYO-mediated tolerance in a dose-dependent manner (**Supplementary, Fig. S8*D***).

**Fig. 4.**
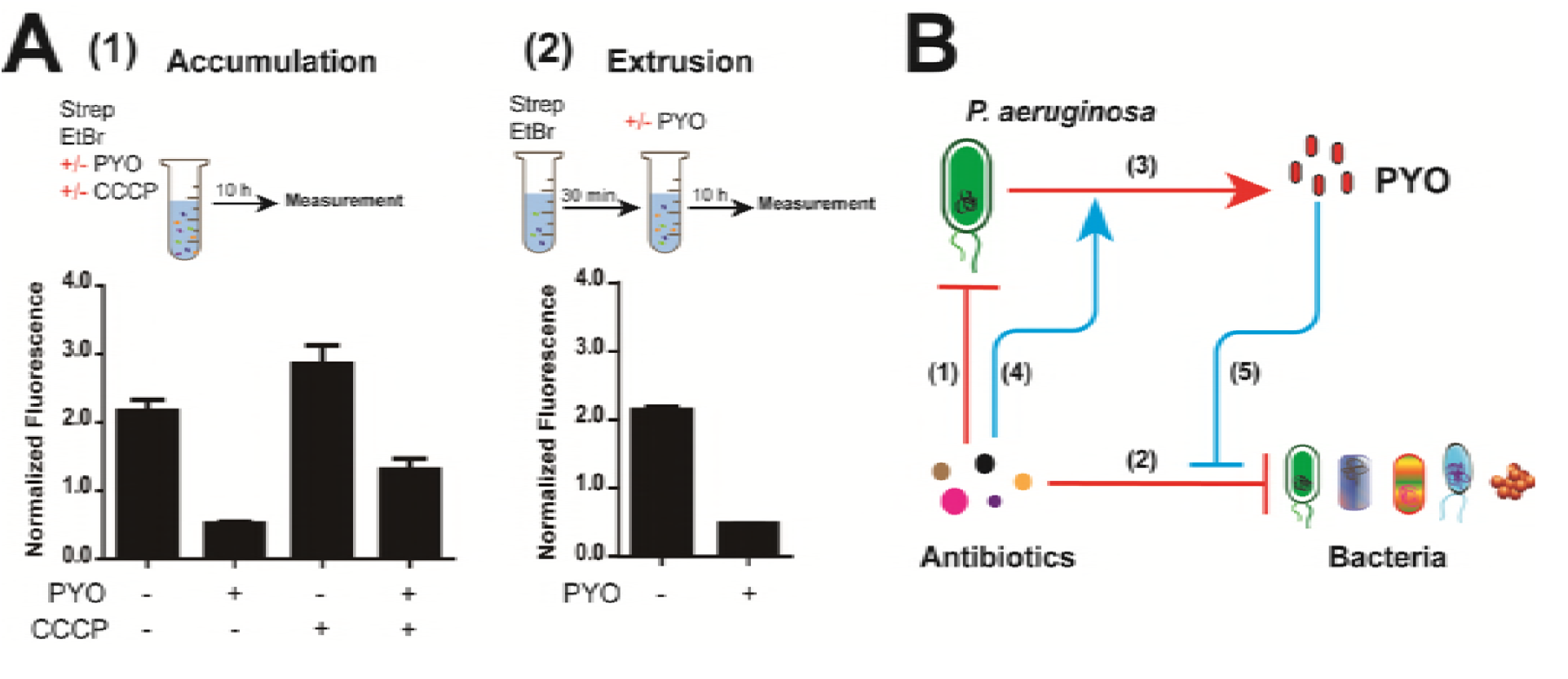
**PYO enhances activity of efflux pumps to export antibiotics**. (**A**) Effect of PYO on EtBr efflux. Accumulation (1) was tested in *P. aeruginosa* PAO1 by the simultaneous addition of ethidium bromide (EtBr), streptomycin (Strep) and PYO. Fluorescence was measured after 10 hrs. Extrusion (2) was tested in *P. aeruginosa* PAO1 pre-treated with streptomycin and EtBr for 30 min, followed by 10 hrs of treatment with PYO. Inhibited pumps could increase the accumulation of EtBr, while activated pumps accelerated the extrusion of EtBr. Fluorescence was normalized to that of streptomycin treated PAO1 in the presence of EtBr. Means ± s.d. were presented throughout (n = 6). (**B**) Model of antibiotics-triggered PYO accumulation to modulate tolerance. Subinhibitory concentrations of antibiotics inhibit bacterial growth (1 and 2), while promoting PYO accumulation in *P. aeruginosa* (3 and 4). PYO at a low level modulates universal antibiotic tolerance for various bacterial species against antibiotics (5).

When the EPs are already expressed at a sufficiently high level, an elevated pump activity in the presence of PYO was primarily due to an enhanced energetic state by the cells, instead of up-regulation of pump genes. For example, various pump gene expression was not regulated (**Supplementary, Fig. S9**). *P. aeruginosa* features a transcriptional factor, SoxR, which is activated by phenazines (20). These results also indicate that the PYO-mediated protection is not due to its regulation of SoxR (**Supplementary, Fig. S10*A***, which can modulate expression of EPs (17). The tolerance was not mediated by SoxR in *P. aeruginosa*, however, this result does not exclude its potential role in other bacteria such as *E. coli*. Moreover, PAO-JP2 strain has multiple quorum sensing (QS) genes knocked out (31), combining the assay with the addition of exogenous QS signals (**Supplementary, Fig. S10*B***), and suggesting a dispensable role of QS in PYO-mediated tolerance. Altogether, we found that the PYO mediated tolerance was neither through working as a signal to regulate the hierarchical QS network (**Supplementary, Fig. S6**), nor through activating SoxR.

## Discussion

The antibiotic-induced early PYO accumulation could result from reduced degradation or increased production of PYO. For example, antibiotics could inhibit enzymes involved in PYO degradation, though such enzymes have not yet been identified in *P. aeruginosa* (51). Alternatively, cells could increase PYO synthesis to counteract stress, akin to that experienced by cells entering the stationary phase. It has been well established that different antibiotics can trigger accumulation of reactive oxygen species (ROS) (52). Likewise, quorum sensing (QS) has been implicated in PYO metabolism (17). However, our experiments demonstrated that the antibiotic-mediated PYO accumulation was unlikely due to these mechanisms (**Supplementary, Fig. S11**).

Collectively, our results reveal a previously unknown mechanism by which *P. aeruginosa* can develop collective antibiotic tolerance (10) mediated by PYO. Both the induction of PYO and the PYO-mediated tolerance are general: all tested antibiotics promoted accumulation of PYO; and PYO conferred tolerance against all antibiotics with the exception of polymyxin B (**Fig. 4*B***).

Moreover, the PYO-mediated protection is not limited to the producing cells; rather, PYO also enhances survival of other *P. aeruginosa* strains and other bacteria. This property highlights a particular challenge in treating infections involving *P. aeruginosa:* incomplete suppression of *P. aeruginosa* by antibiotics would enhance growth of survivors, as well as other pathogens in the growth environment. Nevertheless, the PYO-mediated tolerance results from the cell‘s intrinsic ability to counteract a disruption in the cellular state, whereas PYO itself can be considered as a stress. As such, this tolerance is a double-edged sword. An elevated energy generation can enhance efficacy of certain antibiotics by promoting their uptake (48). Also, if pushed beyond cell’s buffering capacity, the PYO-mediated disruption in the redox balance can inhibit growth (15, 20). Thus, this tolerance mechanism also presents an opportunity for designing treatment strategies that can synergistically enhance effects of antibiotics. For instance, specific inhibitors can be developed to target the multiple enzymes involved for the synthesis of phenazines, antioxidants can be used as adjuvants to counteract the protective effect of PYO.

## Experimental Section

### Chemicals and strains

Antibiotics used in this study include carbenicillin, chloramphenicol, gentamicin, kanamycin, norfloxacin, polymyxin B, streptomycin and tobramycin. Oxidized forms of pyocyanin (PYO) and other phenazines were prepared as previously described (1). Beta-mercaptoethanol, nicotinamide adenine dinucleotide (NADH) and *N*-acetyl-*L*-cysteine (NAC) were used to reduce oxidized form of PYO to its reduced form. 2,6-anthraquinone disulfonate (AQDS) and methylene blue (MB) were used as electron acceptors. Carbonyl cyanide *m*-chlorophenyl hydrazone (CCCP) was used to dissipate the proton motive force. *N,N’*-dicyclohexylcarbodiimide (DCCD) and sodium azide were used to inhibit different subunits of ATP synthase. Ethidium bromide (EtBr) was used to test the function of efflux pumps, and phenylalanine-arginine β-naphthylamide (PAβN) was used to inhibit the efficiency of efflux pumps. *N*-butyryl-*L*-homoserine lactone (C4-HSL) and *N*-3-oxo-dodecanoyl-*L*-homoserine lactone (C12-HSL) were used as quorum sensing molecules. All chemicals were purchased from Sigma-Aldrich and Cayman Chemical.

*P. aeruginosa, Escherichia coli, Salmonella typhimurium, Bacillus cereus, Bacillus subtilis* and *Staphylococcus aureus* strains were used in this study. More details of these strains were shown in **Supporting Information Table S1**.

### PYO accumulation in culture tubes

Different concentrations of antibiotics were added to 4 mL Luria-Bertani (LB) broth (5 g/L yeast extract, 10 g/L tryptone and 10 g/L NaCl, Genesee Scientific) in culture tubes (Genesee Scientific) for *P. aeruginosa* PAO1 and PA14 strains. Cultures at exponential phase were adjusted to A600 nm = 0.2, subsequently 10 μL cultures were resuspended in 4 mL fresh LB medium. All the tubes were cultured in a shaker (New Brunswick Scientific) at 37 °C, 250 rpm. When appropriate, the photos of cultured tubes treated with various antibiotics were taken and cell density (A600 nm) were measured at 600 nm. Excreted PYO was quantified based on the presence of pink to deep red color in acidic solution as described previously (2). Briefly, 2.5 mL supernatant from LB broth was mixed with 1.5 mL chloroform at specified time points. PYO in the chloroform phase was then extracted into 0.5 mL 0.2 mol/L hydrochloric acid (HCl). After centrifugation, the absorbance of the top layer was measured at 520 nm. Concentrations (μg/mL) of PYO were expressed as micrograms of PYO per milliliter supernatant, determining by multiplying the absorbance at 520 nm (A520 nm) by 17.072.

### Growth condition in multi-well plates

For routine growth, bacterial strains were all cultured in 100 μL LB medium in 96-well cell culture plates (Costar, Corning) for long-term measurement with a plate reader (Victor, Perkin Elmer). Exponential growth cultures of bacteria were adjusted to A600 nm = 0.2, then 1:20 diluted in 100 μL fresh LB broth containing various concentrations of antibiotics either in the presence or absence of exogenous 2 μg/mL oxidized PYO or other phenazines. *P. aeruginosa* PAO1 was also cultured in other media, including brain-heart infusion (BHI), M9 and 2xYT media. For growth dynamic measurements, 50 μL mineral oil (Sigma) was added into each well to prevent evaporation. The absorbance was measured by an Infinite 200 Pro plate reader (Tecan) at the wavelength of 600 nm (A600). Furthermore, oxygen permeable sealing membrane (Diversified Biotech) was used to determine whether increasing oxygen concentration could increase antibiotic tolerance. Additionally, LB medium supplemented with different reducing agents (100 μmol/L beta-mercaptoethanol, 15 μmol/L NADH and 13 mmol/L NAC) were simultaneously added with the oxidized form of PYO.

### c-di-GMP assay

*P. aeruginosa* PAO1 was cultivated in LB at 37 °C for 8 h with shaking. 10 mL of cultures (OD600 = 1.2) in the presence or absence of antibiotics at room temperature for 3 hrs, were harvested by quick centrifugation (10 000 g) at 4 °C for 10 min. Cells were washed by 2 mL of water and collected by quick centrifugation (10 000 g) at 4 °C for another 10 min. Cell pellets were re-suspended in 300 μL of water containing 100 µg/L xanthosine 3′5′-cyclic monophosphate (cXMP, Sigma) as the internal standard and incubated at 4 °C for 15 min, then heating at 95 °C for 10 min. After cooling at room temperature for 10 min, the suspension was centrifuged (11 000 g) at 4 °C for another 15 min. Cell pellets were extracted with 200 μL of 65% ice-cold ethanol at 4 °C for twice. The supernatants were evaporated until dryness at 40 °C under nitrogen gas. The residues were re-suspended in 200 μL of 0.5 mmol/L ammonium acetate containing 0.3% (v/v) acetic acid for subsequent UPLC-MS/MS analysis. Briefly, eluent A consisted of 0.5 mmol/L ammonium acetate containing 0.3% (v/v) acetic acid and eluent B was methanol. The injection volume was 10 μL and the flow rate was 0.3 mL/min. Temperature for BEH Shield RP 18 column (Waters) was set at 30 °C. The concentration of c-di-GMP was quantified by Xevo TQ triple quadrupole mass spectrometer (Waters) equipped with an electro spray ionization (ESI) source for multiple reaction monitoring (MRM) analysis in positive ionization mode. The following MRM transitions were detected for the quantification of c-di-GMP at m/z of + 691/152 (quantifier), and confirmatory signals were monitored at m/z of + 691/135 and + 691/248 (qualifier). Final concentration of c-di-GMP was expressed as the ratio of c-di-GMP (ng) to bacterial protein (mg). Protein content was tested by bicinchoninic acid (BCA) assay (Beyotime). Briefly, 2 mL bacterial culture was harvested at 10 000 g and washed by 2 mL of water, then collected by quick centrifugation (10 000 g) at 4 °C for another 10 min. Subsequently, the cell pellet was dissolved in 800 μL of 0.1 mol/L sodium hydroxide and heated at 95 °C for 15 min before testing. The levels of c-di-GMP in *P. aeruginosa* under antibiotic treatments were normalized with that without antibiotics.

### ATP assay

The ATP contents of *P. aeruginosa* PAO1 in LB were measured by the Enhanced ATP Assay Kit (Beyotime, Shanghai, China) according to the manufacture’s instruction. Briefly, 5 mL of early stationary-phase bacteria (A600 nm = 0.7) covered by 1 mL mineral oil in cultural tubes were spiked with 2.5 μg/mL PYO at different time points at room temperature. 4 mL bacterial culture was harvested by centrifugation (10 000 g) at 4 °C for 10 min, then the cell pellets were lysed by 200 μL lysing buffer. Subsequently, the supernatant was obtained by centrifugation (12 000 g) at 4 °C for 5 min. Lastly, the bioluminescence signals were determined by SpectraMax^®^ M5 Microplate Reader (Molecular Devices LLC), based on luciferase <u>catalyzes</u> the formation of light from ATP and luciferin.

### Membrane potential measurement

The membrane potential of *P. aeruginosa* PAO1 was measured by the BacLight^TM^ Bacterial Membrane Potential Kit (B34950, Molecular Probes). Cultures at exponential phase were added to fresh LB medium (approx. 106 CFU) and 100 μmol/L 3,3’-diethyloxacarbocyanine iodide (DiOC_2_(3), membrane stain) with 20 μg/mL streptomycin in a 96-well black plate (Costar, Corning), either in the presence or absence of 2 μg/mL PYO and/or 10 μmol/L CCCP, or both. Subsequently, 50 μL mineral oil was added into each well and the plate was analyzed on an Infinite 200 Pro plate reader (Tecan). The red/green fluorescent intensities for each well were collected with the excitation wavelength at 488 nm and with the emission wavelengths at 525 nm and 590 nm, respectively. The normalized fluorescent values were used to determine the relative changes of proton motive force (PMF).

### Accumulation and extrusion assays

The accumulation of ethidium bromide (EtBr) was carried out in a 96-well black plate (Costar, Corning). Cultures of *P. aeruginosa* PAO1 at exponential phase (A600 nm = 0.2) were 1:20 diluted into 100 μL fresh LB medium, supplemented with 2.5 μg/mL EtBr. 2 μg/mL PYO and 20 μg/mL streptomycin were simultaneously added in the presence or absence of 10 μmol/L CCCP. For the extrusion of EtBr assay, *P. aeruginosa* PAO1 culture was pre-treated with 50 μg/mL streptomycin in the presence of 2.5 μg/mL EtBr for 30 min to allow accumulation of EtBr. The bacteria were centrifuged at 8 000 *g* for 3 min to remove the culture media, and washed by phosphate-buffered saline (PBS, Sigma) twice. The bacterial pellets were replaced by LB broth in the presence or absence of PYO. Lastly, 50 μL mineral oil was added into each well and the plate was measured for 10 h. The absorbance (A600 nm) and fluorescence of the conjugate of EtBr binding to nucleic acid were recorded with an Infinite 200 Pro plate reader (Tecan). The excitation wavelength was set at 488 nm and the emission wavelength was at 590 nm.

### qRT-PCR analysis

The primers used for the quantification of efflux pump gene expression are listed in **SI Appendix. Table S2**. Total RNA was isolated from *P. aeruginosa* cells grown in the presence or absence of streptomycin and PYO with conditions similar to the above. Shortly, overnight cell cultures were 1 to 100 folds diluted into fresh LB medium (pH 7.0) and distributed into a 6-well plate. PYO (2 μg/mL) or/and streptomycin (20 μg/mL) were added and incubated at 30 oC for 2, 5, 10 hrs before the cells were collected for analyses. RNA was extracted using the Qiagen RNAeasy kit (Qiagen) followed by DNase I digestion. cDNA was obtained by reverse transcription of the total RNA using the Applied Biosystems high-capacity cDNA Reverse Transcription kit (ThermoFisher Scientific). qRT-PCR was performed using Power SYBR Green Master Mix (ThermoFisher Scientific) on StepOnePlus instrument (Applied Biosystems) using the housekeeping gene *rpsL* as an internal reference for quantification. Expression of five major RND efflux pump related genes *(oprM, mexB, mexD, mexF* and *mexY)* were assessed using published sets of primers (**SI Appendix. Table S2**). After normalization to the expression of *rpsL*, the levels of five pump genes were compared under different conditions at 5 h, 9 h and 12 h.

### Assays of efflux pumps

Firstly, PAβN as a specific pump inhibitor was used to investigate the efficiency of pumps in the presence of streptomycin. Furthermore, the growth curves of *P. aeruginosa* PAO1 supplemented with 20 μg/mL streptomycin were measured, initially treated with 10 μmol/L CCCP and 1 mmol/L sodium azide, respectively. The growth curves were measured by a plate reader (Tecan) at the wavelength of 600 nm (A600). 50 μL mineral oil was added to prevent evaporation during long term culture.

### Statistical analysis and software

The statistical significance of differences was determined by a Student’s *t*-test. The fluorescent intensities and fold changes in mRNA levels were normalized prior to analysis. GraphPad Prism 5.0 was used for analyzing the data and generating figures.

## Acknowledgments

We thank D. K. Newman (Caltech) for sharing PA14 and its mutants (*Δphz*, *ΔsoxR* and *ΔsoxR-phz*) and purified PYO, PCA, PCN and 1-OHPHZ, H. E. Blackwell (UW-Madison) for sharing PAO-JP2, L. S. Thomashow (WSU) for sharing PAO1-W and its mutants (PAO-mxM and PAO-mxS), D. K. Newman for insightful discussions and critical reading of the earlier drafts of the manuscript, and B. Shao (Beijing CDC) for LC-MS/MS analysis.

## Supporting Information Legends

Material and methods

Tables S1 to S2

Figs. S1 to S10

References (*1-9*)

